# Differential Decay of Multiple eNA Components from a Cetacean

**DOI:** 10.1101/2025.05.09.653189

**Authors:** Pedro FP Brandão-Dias, Megan Shaffer, Gledis Guri, Kim M Parsons, Ryan P Kelly, Elizabeth Andruszkiewicz Allan

**Affiliations:** School of Marine and Environmental Affairs, University of Washington, Seattle, 98105, Washington, USA; Conservation Biology Division, Northwest Fisheries Science Center, National Marine Fisheries Service, National Oceanic and Atmospheric Administration, Seattle, 98112, Washington, USA

**Keywords:** Removal, ddPCR, Bayesian, Experimental, Marine Mammal

## Abstract

Environmental nucleic acids (eNA), such as DNA and RNA, are powerful tools for monitoring biodiversity. Yet, interpreting eNA detections requires understanding of their environmental persistence. We conducted a decay experiment using seawater from an open enclosure to track degradation of six eNA components derived from *Tursiops truncatus*: mitochondrial eDNA of varying lengths, ribosomal eRNA, and messenger eRNA. Targets were quantified over seven days via digital droplet PCR (ddPCR). Decay followed a biphasic exponential model with rapid initial loss (∼24 hours at 15 °C), followed by slower degradation. Cytb messenger eRNA was least stable, disappearing within four hours. Ribosomal eRNA persisted longer but degraded slightly faster than its eDNA counterpart (λ₁ = 0.236 vs. 0.165 hr⁻¹). Longest eDNA fragments decayed more rapidly (λ₁ = 0.190 hr^−1^) than shorter ones (λ₁ = 0.114 hr^−1^). These findings support the use of eDNA fragment length as a proxy for degradation state and reinforce that combining multiple eNA components with distinct stabilities can potentially provide a molecular clock for inferring eNA age. This approach improves the spatiotemporal resolution of eNA-based monitoring, particularly for rare marine-mammal that act as point sources. We also emphasize the importance of explicitly distinguishing between RNA types (ribosomal vs. messenger) in environmental studies, given their divergent stability and interpretability.

## INTRODUCTION

Environmental DNA (eDNA) and, more recently, environmental RNA (eRNA)— collectively referred to as environmental nucleic acids (eNA)—have emerged as powerful tools for ecological monitoring and biodiversity assessments ^1,2^. Unlike traditional direct observation methods, eNA-based surveys rely on indirect detection of species through trace amounts of genetic material that organisms release into their environment ^3^. Due to the eNA accumulation in space, eNA surveys are fundamentally integrative, containing genetic information accumulated over time rather than providing an instantaneous snapshot. This integrative property enhances detection sensitivity compared to nets or visual surveys which may integrate over space but reflect a single point in time; that is, they are snapshot-based methods, making eNA detection analogous to a short movie rather than a single photographic frame. However, the extent to which eNA surveys integrate species presence over both time and space depends on environmental and molecular factors that govern its persistence and movement, thus making the integration variable ^4^.

Accurately interpreting eNA detections requires understanding how much time and space is integrated into each detection event. This, in turn, demands detailed knowledge about the origin, state, transport, and persistence (or decay) of nucleic acids within the environment, collectively referred to as the ecology of eNA ^3,5^. Among these processes, the decay rate of eNAs has been among the most frequently estimated parameters^6–8^. The prominence of decay rate studies stems not only from the relative ease of estimating decay rates compared to transport or production, but also from the fundamental role decay plays in determining the temporal resolution of eNA surveys. While transport primarily governs the spatial resolution of detections ^9^, largely shaped by hydrological, oceanographic, or atmospheric processes, decay dictates how long genetic signals remain detectable, helping set the temporal bounds of the eNA movie, which reflects a steady-state balance among production, transport, and degradation.

Beyond simply establishing an upper temporal limit for eNA detection, certain applications require more precise aging of the biological source of detections within a sample. This is particularly relevant for point sources or rare detections, such as those of marine mammals ^10^ or invasive species ^11^. To address this, previous research has proposed using the ratio of eRNA to eDNA as a “molecular clock” for eNA persistence ^12^. Since eRNA generally degrades faster than eDNA, qualitatively, a sample with a high proportion of eRNA to eDNA suggests a recent biological source, whereas a sample containing only eDNA indicates an older signal ^8,12^. This is analogous to applications in the forensic literature, where the ratio of presence between messenger RNA and DNA, as well as the degradation state of RNA, can be reliably used to infer time since death ^13^.

Nonetheless, this framework extends beyond eDNA/eRNA ratios to any eNA component with distinct degradation rates. Environmental nucleic acids exist in multiple molecular forms, including intracellular, particle-adsorbed, dissolved, nuclear, mitochondrial, and other fractions ^14–16^. Provided these components decay at different rates, then their relative proportions over time and space provide a molecular signature for estimating detection age ^17^. Naturally, this approach can be applied not only to molecular forms of eDNA, but also to experimentally defined fractions such as fragment lengths assessed through multiple genetic markers ^18,19^, variations in eNA particle size distributions captured by sequential filtration ^20,21^, differences between eNA extracted from distinct environmental media, such as sediment versus water ^22^, eDNA to eRNA ratios ^12^, among many others.

In theory, the most effective molecular components for estimating detection age are those with strongly contrasting decay rates, as their relative proportions shift measurably over short timescales, thus enhancing spatiotemporal resolution. However, in practice, the choice of eNA components must balance temporal sensitivity with logistical and methodological feasibility. Numerous studies have measured eNA decay rates, consistently highlighting that these rates are highly context dependent. Decay dynamics vary substantially with environmental conditions such as biological activity and temperature ^23,24^, depend on target organism ^25^, and differ significantly due to methodological differences such as filter pore size and molecular marker length ^17,20,26^. Consequently, generalizing decay rates across systems or conditions is challenging, especially since controlled experimental setups seldom fully represent the complexities of natural environments. As a result, accurate molecular time inference requires decay estimates tailored to or closely approximating the specific environmental conditions and molecular targets of interest.

In freshwater systems, multiple studies have compared eRNA and eDNA decay, generally finding that eRNA—particularly messenger RNA (emRNA)—decays more rapidly than eDNA ^27,28^. In marine environments, however, relatively few studies have directly compared eRNA and eDNA decay. For example, Qian et al.^29^ observed significantly faster decay of prawn-derived emRNA compared to eDNA, particularly under colder temperatures. Conversely, Wood et al.^30^ examined decay rates of worm- and tunicate-derived emRNA, finding only slightly elevated decay rates compared to eDNA. Similarly, Scriver et al. ^31^ found no significant difference in decay between emRNA and eDNA from marine worms. To date, no studies have directly assessed decay rates of ribosomal eRNA (erRNA) in marine environments, although Miyata et al.^32^ noted that erRNA yielded more ecologically relevant metabarcoding detections than eDNA, suggesting greater transience and thus potentially faster decay. Crucially, despite this emerging body of work on eRNA, no studies have yet evaluated the decay rates of eNA components, eDNA or eRNA, specifically derived from marine mammals.

Here, we experimentally assessed the differential decay rates of multiple eNA components from the common bottlenose dolphin, *Tursiops truncatus* (Montagu, 1821). Using seawater sourced from an open environment, netted dolphin enclosure, we conducted a controlled decay experiment, tracking the persistence of eDNA of several molecular lengths, emRNA, and erRNA over time (Fig. 1). We hypothesized that eRNA would degrade faster than eDNA, with emRNA decaying the fastest due to its transient nature. We expected erRNA to persist longer than emRNA due to its structural properties, but still degrade more rapidly than eDNA. Lastly, we anticipated that larger DNA fragments would decay faster than shorter ones, following previous findings ^20,33^ and known fragmentation patterns. By quantifying these decay rates, we aimed to improve the temporal resolution of marine mammal detections and refine our understanding of eNA persistence in marine environments.

**Figure 1:**
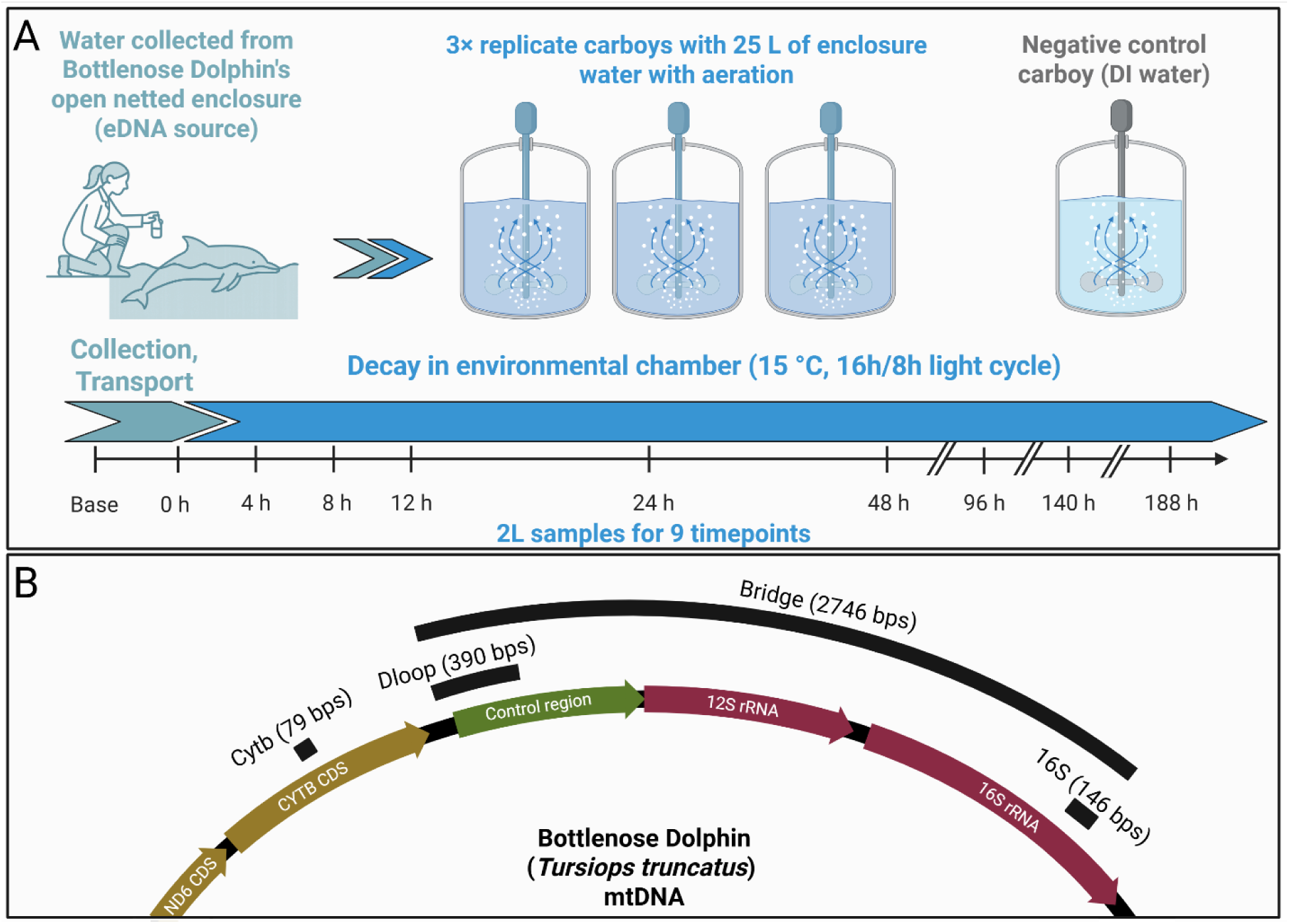
Overview of the experimental design and the four mitochondrial targets quantified. (a) Water-decay experiment. Seawater was scooped from inside an open-net enclosure housing an Atlantic bottlenose dolphin (*Tursiops truncatus*) and poured into three 25 L carboys (biological replicates). A fourth carboy containing de-ionized (DI) water served as cross-contamination control. Carboys were held in a 15 °C environmental chamber with continuous aeration and a 16 h light / 8 h dark cycle. Two-liter subsamples were withdrawn at nine timepoints (0–188 h) for size-fractionated filtration and downstream DNA/RNA extraction. (b) Locations of the mitochondrial assays. Arrows show the relative positions of the 79 bp Cytb amplicon (protein-coding), the 390 bp D-loop amplicon (non-coding control region), and the 146 bp 16 S rRNA amplicon (ribosomal) on the circular dolphin mtDNA. The 2746 bp segment spanning D-loop to 16 S (black arc labelled “Bridge”) represents the minimum intact bridging fragment between 16S and Dloop loci. Figure created in https://BioRender.com.

## METHODS

### Environmental DNA source and decay experiment

A small, managed group of non-native Atlantic bottlenose dolphins (*Tursiops truncatus*) is housed in a defined area along the eastern bank of Hood Canal, Washington, as part of the U.S. Navy’s Marine Mammal Program. These dolphins frequently inhabit netted enclosures adjacent to a pier, where seawater freely exchanges with the surrounding marine environment. In October 2024 we collected water from within one of these netted enclosures containing a single dolphin at the time of sampling. Using a 2-liter pitcher, we filled three replicate 25-liter carboys, which served as the three biological replicates for the decay study (Fig. 1a). To minimize contamination, collectors wore gloves during sampling, and both the pitcher and carboys had been cleaned overnight with 1% bleach and extensively rinsed with deionized water before use.

After collection, water within each carboy was mixed, and a 2-liter baseline sample was filtered from each carboy to assess initial eNA concentrations before transport. These samples were immediately filtered and preserved (see below for filtration and preservation details). The carboys were then sealed and transported at ambient temperature to the laboratory and an environmental chamber for the decay experiment. Temperature loggers placed inside one of the carboys recorded transport temperatures ranging from 16°C to 12°C over the course of ∼3 hours of transport back to the laboratory.

At the laboratory, the three biological replicate carboys were decontaminated externally with bleach before being placed inside an environmental chamber, where they remained for the duration of the experiment. A fourth carboy, filled with deionized (DI) water, was also placed in the environmental chamber as a cross-contamination control. The environmental chamber was maintained at a constant 15°C with a 16-hour daylight cycle throughout the experiment. To ensure water circulation, we inserted air hoses with stones powered by a Whisper 100 (Tetra) into each carboy, in addition to a U-shaped plastic sampling hose, both of which remained in place for the entire experiment. Like the carboys, each of those were cleaned by overnight 1% bleach bath and extensively rinsed with deionized water before use.

To evaluate the decay of eNA over time, 2-liter water samples were collected from each carboy at nine timepoints: 0, 4, 8, 12, 24, 48, 96, 140, and 188 hours (Fig. 1a). At each timepoint, aeration was temporarily turned off to prevent aerosolization, and water was drawn via the plastic sampling hoses. We used sequential filtration to capture eDNA across different particle sizes. Each sample was filtered through a tandem system of three filter housings (Smith-Root, non-self-preserving) equipped with 5 µm, 1.0 µm, and 0.45 µm mixed cellulose ester (MCE) 47 mm filters (Advantec), each backed by polyester drain disks (Sterlitech). Filters were connected downstream to an eDNA Citizen Science Sampler (Smith-Root) for vacuum filtration. Immediately after filtration, filter housings were opened, and sterilized forceps were used to fold each filter twice, sample side in, before placing it into 5 mL Lo-Bind tubes (Eppendorf) containing 2 mL of DNA/RNA Shield (Zymo). Samples were incubated at room temperature for 30 minutes, then frozen at −80°C until DNA extraction within one month.

To prevent cross-contamination, all forceps, filter holders, adapters, and tubing were submerged in 5% bleach baths, thoroughly rinsed with deionized (DI) water, and dried before reuse. Two independent sets of filtration equipment were available, allowing for alternating use between timepoints. Additionally, at the end of each timepoint, we filtered 2 liters of the DI water used for post-bleach rinsing through a 0.45 µm filter as a contamination control to verify the effectiveness of our decontamination procedures.

### DNA and RNA extraction

Nucleotides were extracted from samples using a two-step process. First, frozen filters with buffer were thawed and incubated with agitation at 37 °C for ∼15 minutes to dissolve any precipitation. The samples were then thoroughly vortexed, and the buffer (2 mL) was transferred to an Amicon Ultra-15 30 kDa centrifugal filter unit (Millipore-Sigma) and centrifuged at 4,000 rpm for 30 minutes, concentrating the buffer to ∼400 µL. Both eDNA and eRNA were extracted simultaneously using the Quick DNA/RNA Miniprep Kit (Zymo Research), following the manufacturer’s protocol with a 30-minute proteinase K incubation before adding the binding buffer.

This extraction process results in two sequential extracts (Fig. S2a). The first primarily contains the DNA (hereafter “eDNA”), whereas the second is expected to contain RNA; however, empirical results showed that it also contained a substantial amount of DNA, hereafter referred to as “carryover eDNA”. One-third of this second extract —containing both eRNA and carryover eDNA— was retained for carryover eDNA analysis (see supplement, Figs. S2, S3), while the remaining two-thirds were treated with ezDNAse (ThermoFisher Scientific) at 37 °C for 30 minutes to remove the carryover DNA in preparation for cDNA synthesis.

The DNAse-treated portion of the extract was subjected to a SuperScript IV (ThermoFisher Scientific) first-strand cDNA synthesis reaction using random hexamers in the denaturation step following the manufacturer’s protocol, to convert RNA into cDNA. This product is hereafter referred to as “eRNA.” Each sample was also subjected to a second control reaction, identical in components and cycling conditions, but without the reverse transcriptase enzyme. This reaction product is hereafter referred to as the “No-RT,” and it was used to assess post-DNAse treatment carryover eDNA, ensuring that quantifications from the eRNA extract were from eRNA-derived cDNA rather than residual carryover eDNA. For details on carryover eDNA analysis, see the supplementary material.

### Target eNA quantification

Target eNA were quantified using three assays, all of which target mitochondrial loci (Table 1, Table S1). The first was a 79 bp fragment of *Cytochrome b* (Cytb), a protein-coding gene, from which we quantified three components: eDNA, messenger eRNA (hereafter emRNA), and the No-RT control. The second was a 146 bp fragment of the *16S* ribosomal RNA gene, used to quantify eDNA, ribosomal eRNA (erRNA), and No-RT control. The third was a 390 bp fragment of the *D-loop* control region, from which only eDNA was analyzed as it is a non-coding region.

**Table 1:** Markers analyzed herein. “Bridge” refers to the 16S-Dloop bridge sequence (Figs. 1b, S1).

| Mitochondrial locus | Nucleotide type | Extract | Assay | Target length (base pairs) |
| --- | --- | --- | --- | --- |
| **Cytb** | DNA | eDNA | Cytb monoplex | 79 |
| **Cytb** | mRNA | eRNA | Cytb monoplex | 79 |
| **16S** | DNA | eDNA | 16S-Dloop duplex | 146 |
| **16S** | rRNA | eRNA | 16S monoplex | 146 |
| **Dloop** | DNA | eDNA | 16S-Dloop duplex | 390 |
| **Bridge** | DNA | eDNA | 16S-Dloop duplex* | 2746-16390 |

\* See supplement for calculation details

In addition to these three markers quantified directly by their respective primers and assays, a fourth mitochondrial DNA (mtDNA) component—referred to as the “16S-Dloop bridge sequence”, hereafter “Bridge” for short—was quantified within the duplex ddPCR reaction targeting both *16S* and *D-loop*. This Bridge signal arises from intact (2746+ bps) mtDNA molecules that physically link the two markers (Fig. 1b, see details below and in the supplement). Altogether, this yielded six distinct eNA components to be analyzed: Cytb eDNA, Cytb emRNA, 16S eDNA, 16S erRNA, Dloop eDNA, and Bridge eDNA (Table 1).

All markers were quantified using digital droplet PCR (ddPCR). Cytb eDNA, Cytb emRNA, Cytb No-RT, 16S erRNA and 16S No-RT were quantified using monoplex reactions. Each 22 μL reaction contained 11 μL of ddPCR Supermix for Probes (Bio-Rad Inc., Hercules, CA), 250 nM of probe, 900 nM of each primer (IDT), and 2 μL of template extract. The thermocycling conditions were: 4°C for 10 min; 95°C for 10 min; 45 cycles of 94°C for 30 s and 60°C for 60 s (annealing/extension); followed by 98°C for 10 min and a final hold at 4°C.

For *16S* and *D-loop* eDNA, we used a duplex ddPCR reaction. Each 22 μL reaction included 11 μL of Supermix, 900 nM of *D-loop* primers, 600 nM of *16S* primers, 250 nM of each probe, and 2 μL of extract. Given the longer amplicon sizes, we adjusted cycling conditions: 4°C for 10 min; 95°C for 10 min; 44 cycles of 94°C for 30 s, 56°C for 30 s (annealing), and 72°C for 120 s (extension); followed by 98°C for 10 min and a 4°C hold. This duplex design allowed us to simultaneously quantify 16S, D-loop, and the mtDNA bridge between them (Fig. 1b).

As mentioned above, this duplex reaction also allowed us to quantify a fourth marker, Bridge, given by ddPCR droplet sorting and linkage between markers (Fig. S1). Droplet PCR partitions the reaction into ∼20,000 droplets, enabling absolute quantification of target molecules using Poisson statistics. If targets are unlinked, they sort into droplets independently, and double-positive droplets arise randomly. However, due to the circular nature of mtDNA, *16S* and *D-loop* are physically linked in intact molecules. The shortest span between them is 2746 bp, though linkage could span the full ∼16,388 bp mitochondrial genome (Louis et al., 2023; Xiong et al., 2009). Thus, in a hypothetical situation where all mtDNA molecules are intact in the sample, all droplets containing 16S would automatically also contain Dloop, and vice versa, making all droplets positive for both markers.

However, because eDNA in environmental samples is partially fragmented, droplets in the duplex assay include a mixture of single-positive and double-positive events. Double-positive droplets can arise from two mechanisms: (1) random co-localization of independently degraded DNA fragments containing 16S and D-loop sequences, or (2) intact DNA molecules physically bridging the two markers (i.e., ≥2746 bp). Using the counts of single-positive droplets for each marker, one can calculate the expected number of double-positive droplets that would result from mechanism (1) alone. This expected value is then subtracted from the observed number of double-positive droplets, and the remainder is attributed to mechanism (2), providing an estimate of the number of intact, bridged molecules. See supplements for full derivation and schematics.

For all ddPCR assays, droplets were generated using the AutoDG Droplet Generator (Bio-Rad Inc.), thermocycling was performed on C1000 Touch Thermal Cyclers (Bio-Rad Inc.), and droplet fluorescence was read using a QX200 Droplet Reader (Bio-Rad Inc.). Each plate included one positive control and three no-template controls (NTCs), used to define positive thresholds and screen for contamination.

Finally, to confirm the absence of non-target marine mammals in the decay experiment, we opportunistically performed metabarcoding on eDNA samples filtered at the first timepoint of each carboy. We detected no other cetacean species in the decay water (Tables S2, S3). Details of metabarcoding methods and results are provided in the Supplement.

### Statistical analysis and decay rate estimation

We built a Bayesian hierarchical model to jointly estimate the eNA concentrations and the decay rates for each component and marker combination *i* (Table 1) from the observed ddPCR droplet counts (W). To estimate the eNA concentrations from ddPCR droplet reader observations, we followed the same approach used in Guri et al. ^36^ where the number of positive droplets from technical replicates (*r*) of the same sample collected at time (*t*) are modeled as a Binomial distribution:

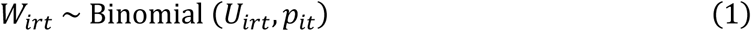

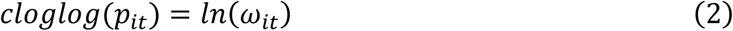

Where *p*_*it*_ is the probability that an individual droplet positively amplifying the target locus *i* at sampled time *t* from the total number of droplets generated *U_irt_* and *ω_it_* is the respective eNA molecules (in copies/μL reaction volume)

For inferring the eNA concentration in the mesocosm (*C_i_*; units copies/L) we normalize for the volume filtered, eluted, and diluted as follows:

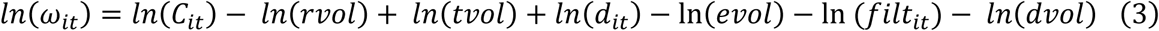

where *rvol* is the total ddPCR reaction volume (22 µL), *tvol* is the template volume added to the reaction (2 µL), *d*_*i*_ is a sample-specific template dilution (caused by reverse transcription reaction), *evol* is the extraction eluted volume (50 µL), *filt*_*i*_is the sample-specific filtered volume, and *dvol* is the ddPCR droplet volume (∼0.85 nL), as specific by Bio-Rad and used by Guri et al. (2024).

To calculate decay for each of the components (c), we tested six different decay models and ultimately used the Biphasic Exponential Decay model which yielded the highest likelihood (see Supplements for model testing). The biphasic exponential decay model assumes two exponential decay phases with a transition time specific for each component and marker (t_ix_):

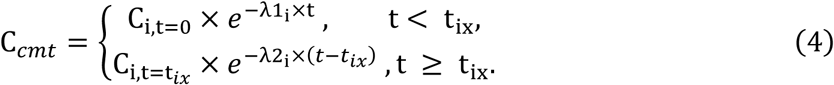

where C_*it*_ is the eNA concentration of each component (*c*) and marker (*m*) at time (*t*); C_i,t=0_ is the respective initial concentration at t = 0; λ1_i_ and λ2_i_ are the decay rate constants for the first and second phases, respectively; t_ix_ is the time where decay rate changes; and 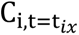 is the concentration at t_ix_ determined as 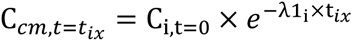.

The concentrations C_*cmt*_ are jointly modeled from two technical replicates (r), and the decay rates (λ1_i_ and λ2_i_) and t_x_ for each component and marker combination considering all three biological replicates. The model was implemented in the Stan language with the R package Rstan ^37^ running four independent Markov chain Monte Carlo (MCMC) chains with 5000 warmup and 10000 sampling iterations. Model’s effective sample size was above 500, and model’s *R̂* convergence parameter was < 1.005 for all estimated parameters.

## RESULTS

### Controls

All target eNA assays from the fourth (control) carboy, filtration negative controls, extraction blanks, and no reverse transcriptase (No-RT) controls yielded zero positive droplets, with the exception of occasional single positive droplets in No-RT controls. These rare events indicate minimal residual DNA carryover in RNA extracts despite DNase treatment. To account for this, we subtracted the droplet count observed in each No-RT control from the corresponding eRNA droplet count of the same sample before proceeding with downstream analyses.

### Little eDNA found in smaller pore sizes

To assess the particle-size distribution of dolphin eDNA, we filtered each time-point sample through a 3-stage serial filtration system (5 µm, 1.0 µm, 0.45 µm). Preliminary sample screening of the first timepoint samples found that >95% of the Cytb signal was captured on the 5 µm filter at time zero. Therefore, all downstream analyses used this fraction exclusively (Fig. 2).

**Figure 2:**
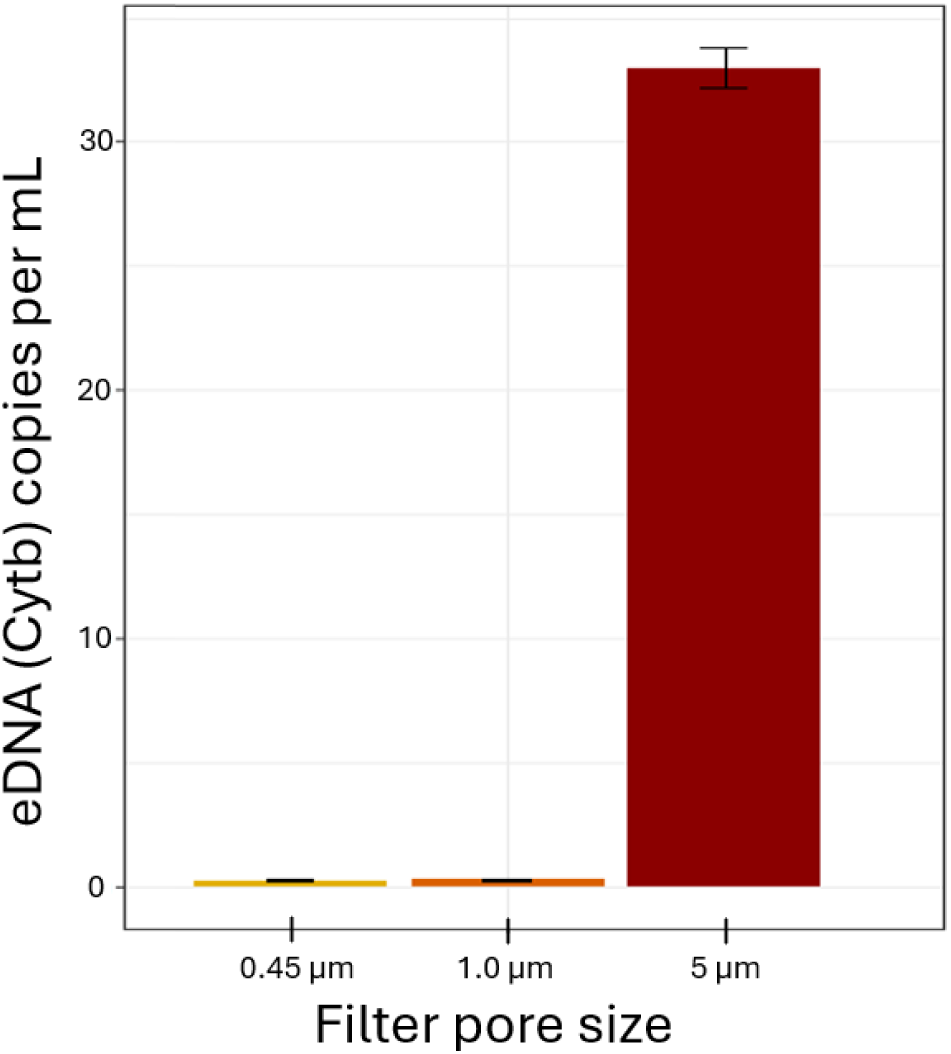
Particle size distribution of Bottlenose Dolphin eDNA in the first timepoint. Concentration of dolphin mitochondrial Cytb eDNA recovered at the first sampling time-point as a function of pore size. Filters with pore sizes < 5 µm retained negligible eDNA, so subsequent analyses rely exclusively on extracts from the 5 µm fraction.

### Cytb messenger RNA was short-lived, and eNA decay was biphasic

We quantified decay of six different eNA components, including four eDNA markers of varying length (Fig. 1b), one erRNA marker, and one emRNA marker (Table 1).

We detected dolphin-derived eDNA and 16S ribosomal eRNA across multiple timepoints, with signals persisting up to 48 hours after the start of the experiment. In contrast, Cytochrome b (Cytb) messenger eRNA was only detected in the first pre-transport sample (Figs. 3, S4). This signal was lost during the ∼3-hour transport period, and no Cytb emRNA was detected in subsequent laboratory samples, suggesting extremely rapid degradation.

**Figure 3:**
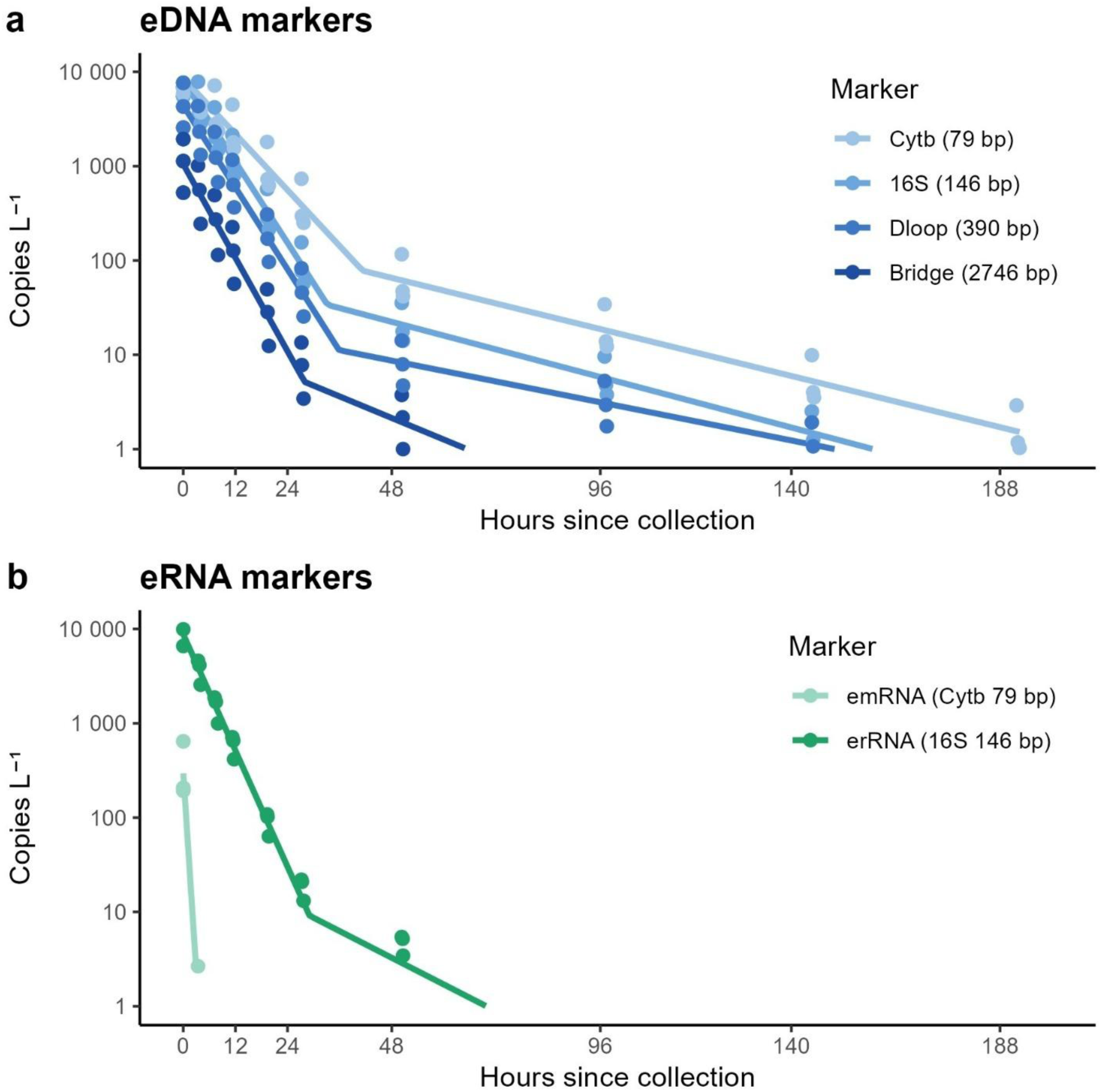
Biphasic-model decay of dolphin environmental nucleic acids (eNAs). (a) Mitochondrial DNA targets. (b) RNA targets. Solid lines are the posterior mean trajectories from the biphasic Bayesian ddPCR model; filled circles are the model-predicted concentrations at each sampling time for every biological replicate (three carboys). Colors distinguish markers and correspond to amplicon length: Cyt b 79 bp, 16 S 146 bp, DLL1 390 bp, and the 2 746 bp Bridge (DNA); Cyt b emRNA 79 bp and 16 S erRNA 146 bp (RNA). Axes are log-scaled; concentrations are expressed as copies L⁻¹.

Beyond this specific marker behavior, we observed that environmental nucleic acids (eNA) generally followed a biphasic decay pattern. This was especially clear for short eDNA fragments, which showed a steep decline within the first 48 hours, followed by a plateau phase in which low concentrations persisted through the end of the 7-day experiment. To formalize this observation, we compared six candidate decay models (Supplement), and the biphasic exponential model provided the best fit to the data based on leave-one-out cross-validation (Table S3).

Decay rate estimates varied by marker (Table 2). Although Cytb emRNA showed the fastest apparent initial decay (λ₁ = 1.615 hr⁻¹), this estimate is based on a single detection and subsequent non-detections, and should be interpreted with caution. Among consistently detected markers, 16S erRNA had the highest initial decay rate (λ₁ = 0.236 hr⁻¹; Fig. 3b).

**Table 2:** Posterior decay rates estimated across markers. “λ_1_” is the decay rate for the first phase of the exponential decay; “λ_1_ 2.5%” and “λ_1_ 97.5%” represent it’s 95% confidence interval; “t_x_” is the time break at which the decay rate was found to change from the first phase to the second; and “λ_2_” is the decay rate for the second phase of the exponential decay. NA values indicate no detections were made at the relevant timepoint, making values unidentifiable.

| Marker | $\lambda_1$ | $\lambda_1$ 2.5% | $\lambda_1$ 97.5% | $t_x$ | $\lambda_2$ |
| --- | --- | --- | --- | --- | --- |
| **Cytb eDNA** | 0.114 hr<sup>-1</sup> | 0.110 hr<sup>-1</sup> | 0.118 hr<sup>-1</sup> | 41 hrs | 0.026 hr<sup>-1</sup> |
| **Cytb emRNA** | 1.615 hr<sup>-1</sup> | 0.877 hr<sup>-1</sup> | 2.763 hr<sup>-1</sup> | NA | NA |
| **16S eDNA** | 0.165 hr<sup>-1</sup> | 0.161 hr<sup>-1</sup> | 0.169 hr<sup>-1</sup> | 33 hrs | 0.028 hr<sup>-1</sup> |
| **16S erRNA** | 0.236 hr<sup>-1</sup> | 0.207 hr<sup>-1</sup> | 0.267 hr<sup>-1</sup> | 29 hrs | 0.054 hr<sup>-1</sup> |
| **Dloop eDNA** | 0.166 hr<sup>-1</sup> | 0.160 hr<sup>-1</sup> | 0.173 hr<sup>-1</sup> | 36 hrs | 0.021 hr<sup>-1</sup> |
| **Bridge eDNA** | 0.190 hr<sup>-1</sup> | 0.175 hr<sup>-1</sup> | 0.206 hr<sup>-1</sup> | 28 hrs | 0.044 hr<sup>-1</sup> |

For eDNA, molecular length was a strong predictor of persistence (Fig. 3a). Shorter targets, such as Cytb (λ₁ = 0.114 hr⁻¹), decayed more slowly, while longer targets like the Bridge fragment spanning 16S and D-loop decayed faster (λ₁ = 0.190 hr⁻¹). The transition between the two decay phases consistently fell between 24 and 48 hours, and all targets excibitted a slower decay rate on the second phase (λ_2_ between 0.054 and 0.021, Table 2).

## DISCUSSION

This study provides the first empirical estimates of decay rates for multiple environmental nucleic acid (eNA) components derived from a marine mammal. Using a controlled experiment with seawater from a dolphin enclosure, we documented the degradation of mitochondrial eDNA fragments of different lengths, ribosomal eRNA, and messenger eRNA from *Tursiops truncatus* over seven days. We found that decay patterns were highly dependent on molecule type and length, with messenger RNA degrading the fastest, ribosomal RNA decaying slightly faster than its DNA counterpart, and longer DNA fragments decaying more rapidly than shorter ones. Across most targets, decay followed a biphasic trajectory, with a steep initial decline and a slower second phase. These results confirm that eNA components differ markedly in environmental stability and underscore the potential of combining multiple markers to improve the temporal resolution of eNA-based detections.

### Rethinking molecular strategies for eDNA age estimation

The ratio between eDNA and emRNA has been proposed as a molecular proxy for time since eDNA shedding^12^, consistent with the spatiotemporally variable eDNA composition hypothesis ^17^. This approach assumes that RNA decays more rapidly than DNA, such that high RNA:DNA ratios indicate recent activity. However, our results suggest that messenger RNA (emRNA) is not well-suited for this application in marine systems. Cytb emRNA was only detectable at the initial timepoint and degraded beyond detection during transport—highlighting the molecule’s extreme lability. Similar challenges have been noted in other marine studies, where emRNA is often either undetectable or not measurably different in decay from eDNA ^30,38^. Beyond this, emRNA analysis requires transcript-specific assays, rapid stabilization, and higher-cost laboratory workflows ^2,39^, which limit its practicality for most field applications.

Instead, our findings support the use of eDNA fragment length as a proxy for degradation state. Longer mitochondrial markers decayed consistently faster than shorter ones, as shown here and in prior work ^20,33^, making them well-suited for relative age estimation ^19^. This method is logistically simple—relying on standard eDNA workflows and multiplexed assays—and broadly applicable across taxa and systems. Importantly, quantitative estimation of eDNA age requires both accurate decay rates and knowledge of the initial proportion of each component, so that their relative abundance over time can be meaningfully interpreted. When targeting mitochondrial loci, the physical linkage of gene regions provides a useful simplification: short and long fragments are expected to originate at roughly equal copy number, enabling decay-based inferences without requiring independent normalization of starting ratios. On the other hand, because mRNA expression levels change by context, the starting proportion between mRNA and DNA in environmental samples will always vary, and initial proportions cannot be accurately established. Thus, pairing mitochondrial markers of different lengths potentially offers a scalable, cost-effective alternative to RNA-based strategies. Rather than relying on a single target, a duplex assay can be designed to simultaneously amplify two mitochondrial regions with a deliberate difference in length, maximizing the contrast in decay rates while maintaining amplification efficiency.

### Mechanisms and practical significance of Biphasic decay

Dolphin eDNA in our experiment followed a biphasic exponential decay: a steep first-phase loss lasting ∼48 h, followed by a low-level “tail” that persisted for days. Similar two-phase kinetics have been reported from rivers, lakes, and coastal waters ^25,40,41^, but often go unrecognized when decay experiments end after only one or two days ^42,43^. We detected no compelling biphasic signal for either ribosomal or messenger eRNA, but it is possible RNA also follows the same pattern.

The processes that generate two-phase decay remain unresolved. Three non-exclusive explanations have been previously proposed: (i) physical shielding, in which DNA adsorbed to mineral or organic particles become inaccessible to nucleases ^44,45^; (ii) encapsulation, whereby intact cells and mitochondria only release DNA after membrane rupture ^46,47^; and (iii) particle-size-dependent removal, where large aggregates settle or are grazed quickly while finer particles or dissolved DNA persist in suspension ^47^. Given these mechanisms are associated with different eDNA states, disentangling them will require experiments that characterize or separate DNA states.

Nonetheless, from a monitoring perspective, the slow second phase probably contributes little to routine eDNA surveys: most metabarcoding data will be dominated by the highly concentrated but rapidly decaying fraction, as the residual tail is easily swamped by fresh inputs. Nevertheless, understanding the process could improve source-age estimates, especially when low concentrations are observed.

### Distinguishing Ribosomal and Messenger eRNA in Environmental Studies

In our study, we found cytochrome b (Cytb) messenger eRNA (emRNA) at very low concentrations, detectable only at the initial time point before water samples were transferred to the environmental chamber. Low levels of Cytb emRNA in environmental samples has been found in other studies ^48^, and these low concentrations are the likely explained by low generation and expression of Cytb from eNA-generating tissue. In contrast, ribosomal eRNA (erRNA) was present at much higher concentrations in our samples, consistent with its constitutive expression, and decayed only slightly faster than its equivalent DNA. This contrast in stability and concentrations of both forms of RNA aligns with well-documented differences in RNA stability from forensic and molecular biology research, where ribosomal RNA’s secondary structure may protect it from degradation ^49,50^. These findings highlight a fundamental distinction between erRNA and emRNA: while erRNA in found in high concentrations and can be reliably detected, emRNA is highly transient, making it a good candidate for detection of more recent targets, but potentially unreliable in yielding consistent taxa detections given its low starting concentrations.

The rapid degradation of mRNA is associated with its biological function. Within cells, mRNA is a highly labile molecule that allows for rapid and dynamic gene expression regulation ^51^. To maintain precise control over protein synthesis, cells possess multiple pathways for degrading mRNA efficiently, including exonuclease-mediated decay and other complex biomolecular pathways ^52,53^. Additionally, mRNA’s single-stranded structure and chemical composition (i.e., an extra hydroxyl group) make it inherently less stable than DNA, leading to its rapid degradation both within cells and in the environment, particularly in high temperatures and alkaline conditions ^27,54^. In contrast, rRNA is constitutively (permanently) expressed in large quantities, representing over 80% of all RNA within cells in many estimates, and plays a structural role in ribosome assembly ^55–57^. Its highly structured secondary and tertiary conformations enhance its resistance to enzymatic degradation ^50,58^, which likely extends to higher stability in the environment as well.

Despite these well-established molecular differences, the term “eRNA” is often used interchangeably in the eDNA literature to refer to both erRNA and emRNA, despite their fundamentally different properties. This conflation can lead to incorrect assumptions about eRNA persistence, decay rates, and ecological interpretability. Specifically, our results and previous literature indicate that emRNA degrades much faster than eDNA in environmental samples, and its initial concentration depends heavily on gene expression in the tissues shedding eDNA. Because gene expression varies across tissues, developmental stages, and environmental conditions, interpreting emRNA abundance requires transcriptomic knowledge of the target species—information that is often unavailable, but can be used to inform emRNA assay development ^39^. Consequently, assays designed for both eDNA and emRNA detection (e.g., targeting Cytb, as tested here) may not be widely effective, as the emRNA marker will have to be customized for each application.

In contrast, erRNA is more abundant, making it a more stable biomarker but unsuitable for most other applications where emRNA would be advantageous over eDNA, such as metabolic inferences ^2^. Supporting this, previous metabarcoding studies using erRNA and eDNA to detect species with a 12S marker found minimal differences in detection rates, though erRNA showed faster species accumulation curves ^38,59^. Macher et al.^38^ also tested emRNA-based detection using cytochrome oxidase I (COI), and as expected, emRNA was less effective than eDNA in species detection due to its rapid degradation (see above). However, its faster decay resulted in stronger spatiotemporal patterns, reflecting its short-lived nature in the environment.

These findings reinforce the importance of distinguishing rRNA from mRNA in environmental studies. The ecological patterns observed for rRNA cannot be assumed to apply to mRNA, and vice versa. Given their fundamental biological and functional differences, treating both molecules collectively as “eRNA” is misleading and can obscure key differences in their degradation rates and interpretability in environmental monitoring. Future research should clarify this distinction and ensure that eRNA studies account for the vastly different behaviors of these two RNA types in environmental contexts.

### Dolphin eDNA Predominantly Retained in Large Particle Fractions

Interestingly, we found that the vast majority of dolphin eDNA was captured on the largest pore size filter (5 µm), with only residual amounts recovered in smaller size fractions (Fig. 2). This pattern contrasts with previous studies on the particle size distribution (PSD) of eDNA from other marine taxa, such as teleost fish (Jo et al., 2019) and elasmobranchs ^60^, whose eDNA is more evenly distributed down to small filter pore sizes (< 1 µm). While eDNA PSD is known to vary with environmental conditions and organism ^44,61^, the pattern observed here closely resembles that reported for freshwater fish in the presence of clay or titanium dioxide, which strongly adsorb to eDNA ^62^. However, such particles were not present in our samples in substantial quantities. There are two possible alternative explanations for the observed PSD. First, it is possible that filtration failure occurred at the smaller pore sizes, preventing accurate capture of eDNA in those fractions. Nonetheless, we observed the same pattern across three replicate carboys, which were filtered separately. Alternatively, if experimental error can be excluded, the results may indicate that marine mammal eNA is predominantly associated with larger particles—possibly due to tissue origin, mucus binding, or aggregation. If this is the case, it has important implications for sampling design: using larger pore size filters may enhance eDNA yield and improve detection probability for marine mammals by allowing greater volumes to be processed.

### Carryover DNA and components of eDNA

Commercially available extractions columns were originally designed to extract high molecular yield DNA from tissue samples. However, they have been commonly used for extracting eDNA from filters over the past two decades and have yielded sufficient DNA for most applications ^1^. To our knowledge, no other studies have captured and quantified eDNA that passes through the DNA extraction columns to quantify this inefficiency. Here, we found a substantial amount of eDNA passed through the initial column, largely comprising shorter DNA molecules as measured by a TapeStation (Supplement Fig. 2). This finding points to a potential blind spot in many eDNA workflows: studies seeking rare targets or high detection sensitivity should be cautious about relying on column-based extraction methods, despite their convenience and scalability.

In addition to quantifying the amount of carryover eDNA, we also quantified the decay rate of carryover eDNA with one marker (Cytb) and found it to decay significantly slower than the eDNA fraction captured on the first extraction column (Fig. S3). This is likely due to a cascading effect, where the decay of longer eDNA molecules creates shorter eDNA molecules, resulting is a slower apparent decay of the smaller fragment size ^17^. Nonetheless, it is unclear if this method of splitting eNA components (i.e., eDNA vs. carryover eDNA) will be useful in the future because the fraction of eNA passing through the first column may not be stable, and may vary substantially under varying total quantities of DNA in the sample and other chemical characteristics of the sample. Thus, while splitting eNA by molecule length (in basepairs) may find its applications, separation of multiple eNA components may be more reliable with other methods such as sequential filtration ^20,63^, evaluating multiple markers of varying lengths as here, or differential extraction methods ^14^.

## ACKNOWLEDGEMENTS

The authors thank the U.S. Naval Base Kitsap-Bangor for logistic support, Kate Bertko for administrative and field support, Krista Nichols, Amy Van Cise, Elizabeth Brasseale, and Olivia Scott for field sampling support, and Andrew Olaf Shelton for sampling support and assistance discussing the STAN decay model. This research was funded by the Office of Naval Research under Award Number (N00014-22-1-2719) and supported through collaborative research efforts with the U.S. Navy’s Marine Mammals Program. Any use of trade, firm, or product names is for descriptive purposes only and does not imply endorsement by the U.S. Government.

## AUTHOR CONTRIBUTIONS

All authors worked together to conceive and design the experiment and analyses; MS, KP, RK, EAA planned fieldwork and sampled water; PBD prepared and executed decay experiment; MS designed assays; PBD performed lab work; PBD and GG wrote the hierarchical Bayesian model; PBD, GG and EAA analyzed and visualized data; KP, RK, EAA obtained funding and supervised the project. All authors discussed results, contributed to and approved the final manuscript draft.

## DATA AVAILABILITY STATEMENT

Raw dataset and corresponding metadata will be available as supplement files upon publication

## COMPETING INTERESTS

The authors declare no competing interests.

## SUPPORTING INFORMATION

Additional description of ddPCR methods, additional visualizations of the raw data, and analysis of carryover eDNA is provided in a pdf as Supplementary Information.

## Notes

### Competing Interest Statement

The authors have declared no competing interest.

### Summary of Updates

Changes were added in accordance with NOAA internal review. In addition, a bug was found in the decay rate code, which resulted in a change in the values of decay rates (the results/ interpretation is the same, the values are simply different magnitudes).

