## Supporting Information for "Differential Decay of Multiple eNA Components from a Cetacean"

### Estimating droplet counts from intact mitochondrial molecules in duplex ddPCR

As described in the main text, duplex droplet digital PCR (ddPCR) enables simultaneous quantification of two mitochondrial markers—in this case, Tt-DLoop and Tt-16S—by partitioning each reaction into approximately 20,000 nanoliter-scale droplets. Each droplet is scored as negative, single-positive (for either DLoop or 16S), or double-positive (for both markers), based on fluorescence amplitude in two channels (FAM and HEX). Because mitochondrial DNA (mtDNA) is circular, the two target loci are physically linked in intact molecules. The shortest possible linear distance between the DLoop and 16S primer binding sites is 2,746 bp, though intact molecules may span the full mitochondrial genome (~16.4 kb; Louis et al., 2023; Xiong et al., 2009). Accordingly, the presence of both markers in a droplet may indicate either (1) independent DNA fragments containing each amplicon co-localizing randomly, or (2) a single long fragment that bridges both targets (Figure S1).

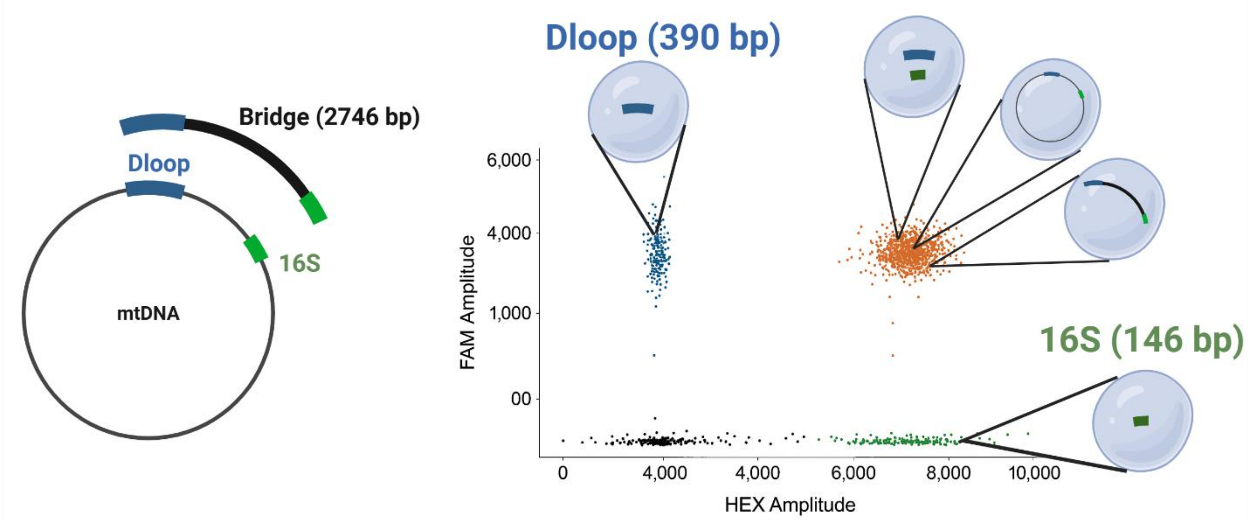

**Figure S1: Illustration of duplex ddPCR-based inference of intact mitochondrial DNA fragments.** Left: Diagram of mitochondrial DNA showing primer binding sites for Tt-DLoop (blue) and Tt-16S (green), and the ~2.7 kb "bridge" segment linking them. Right: Scatterplot of droplet fluorescence (FAM vs. HEX amplitudes) from a hypothetical duplex ddPCR reaction. Droplets are classified into four categories: double negative (black), single positives for DLoop (blue) or 16S (green), and double positives (orange). Insets depict example droplet contents: single-target fragments vs. a long intact DNA molecule spanning both amplicons. Double-positive droplets exceeding the expected co-occupancy rate are attributed to these long fragments. Created in https://BioRender.com.

**Table S1: Oligonucleotide sequences for the assays used in this study.**

| Mitochondrial locus | **Oligo Name** | **Role** | **Sequence** | **Reference** |
|---|---|---|---|---|
| **Cytochrome b (Cytb)** | Ttru-cytb_F | Forward | TTATTCTTCCATTCATCATCAC | Xiong et al., 2025 |
| | Ttru-cytb_R | Reverse | GTGGGGTTGTTGGATCCTGT | Xiong et al., 2025 |
| | Ttru-cytb_P | Probe | /5FAM/AATAGTAGG/ZEN/TGAACGGCTGCCA/3IABkFQ/ | This study |
| **16S** | Ttru-16S_F | Forward | AGGGTTTTACTGTCTCTTACTCTT | This study |
| | Ttru-16S_R | Reverse | CTTTGTTATCCCTTGGTGGTATTA | This study |
| | Ttru-16S_P | Probe | /5-HEX/ATAATGCAA/ZEN/TAAGACGAGAAGACCCTATGG/3IABkFQ/ | This study |
| **Dloop (Control region)** | DL1-HL_F | Forward | CACCCAAAGCTGRARTTCTDYATAAACT | This study |
| | Oordlp4 | Reverse | GCGGGTTGCTGGTTTCACG | Baker et al., 2018 |
| | Ttru-DL_P | Probe | /5FAM/ACTGACGTA/ZEN/GTACTGTGATGTTGTGGTAACTGTAC/3IABkFQ/ | This study |

We aim to isolate and quantify the second mechanism—droplets containing intact mtDNA fragments ≥2,746 bp. To do so, we first estimate the number of double-positive droplets expected by random co-localization, and then subtract that value from the observed double-positive count. The remainder is attributed to physically intact mtDNA templates.

Let N denote the total number of accepted droplets; $n_A$ and $n_B$ be the number of droplets positive for *Tt-16S* and *Tt-DLoop*, respectively (each including double-positive droplets); and let $n_{AB}^{obs}$ represent the number of observed double-positive droplets.

Assuming that fragments containing Tt-16S and Tt-DLoop enter droplets independently, the probability of both appearing in the same droplet is the product of their individual positive fractions:

$$p_{A=}\frac{n_A}{N}, \quad p_{B=}\frac{n_B}{N}$$

The expected number of double-positive droplets under random co-localization is therefore:

$$E\left[n_{AB}^{rand}\right] = Np_A p_B = \frac{n_A n_B}{N}$$

This is the number of droplets we would expect to fall into the double-positive quadrant of the fluorescence plot by chance if no intact bridge molecules were present.

Then, we define the number of double-positive droplets attributable to *intact* molecules as the difference between the observed and expected values:

$$n_{AB}^{bridge} = n_{AB}^{obs} - \frac{n_A n_B}{N}$$

Finally, since this calculation can occasionally yield negative values due to sampling variance, we enforce a non-negativity constraint, making the minimal number of droplets to be zero.

This final calculated value, $n_{AB}^{bridge}$, represents the number of droplets that likely contain a single DNA fragment spanning both the DLoop and 16S regions—i.e., a molecule of at least 2,746 bp in length. These counts are carried forward into the Stan model to infer the decay dynamics of intact mtDNA over time.

While not applied in this study, future experiments could incorporate selective restriction enzymes that cleave only one side of the bridge region, enabling more precise discrimination between linked and unlinked fragments.

### Coextraction Carryover analysis

To assess the performance of our DNA/RNA coextraction protocol and characterize the persistence of DNA contaminants in RNA extracts, we implemented a two-step spin column workflow (Figure S2A). As described in the Methods, environmental DNA (eDNA) is first recovered from the initial eluate after column binding, while RNA is recovered from the second eluate after ethanol precipitation of the flowthrough. However, fragment analysis (Figure S2B) revealed substantial residual DNA in this second eluate—hereafter referred to as "carryover eDNA"—indicating incomplete separation. TapeStation results show that while eDNA isolated from the first eluate consists of high–molecular-weight fragments, the carryover DNA is systematically shorter.

These findings underscore two major considerations. First, DNase treatment of RNA extracts is essential to avoid misquantification of eRNA due to carryover eDNA contamination. This justifies the inclusion of No-RT controls across all analyses. Second, they raise broader concerns about the suitability of silica column–based methods for eDNA research. These methods are optimized for tissue-derived DNA and preferentially recover high–molecular-weight fragments, leading to loss of smaller eDNA fragments that may be critical for detecting degraded or low-abundance signals.

To investigate how carryover DNA behaves over time, we modeled its degradation dynamics in comparison to the main eDNA fraction for a mitochondrial Cytb marker (Figure S3). Both fractions followed biphasic exponential decay, but with notable differences. The initial decay rate ($\lambda_1$) was lower for carryover DNA, and the secondary decay rate ($\lambda_2$) was higher, resulting in a shorter overall transition time (tx). These patterns suggest that carryover eDNA may include more stable or protected DNA molecules, or may accumulate shorter fragments over time as larger ones degrade and bypass the initial column step. This "cascade effect" (Brandão-Dias et al., 2023b) could confound estimates of decay rates and persistence, especially when RNA-derived cDNA is used as a proxy for activity or recent presence.

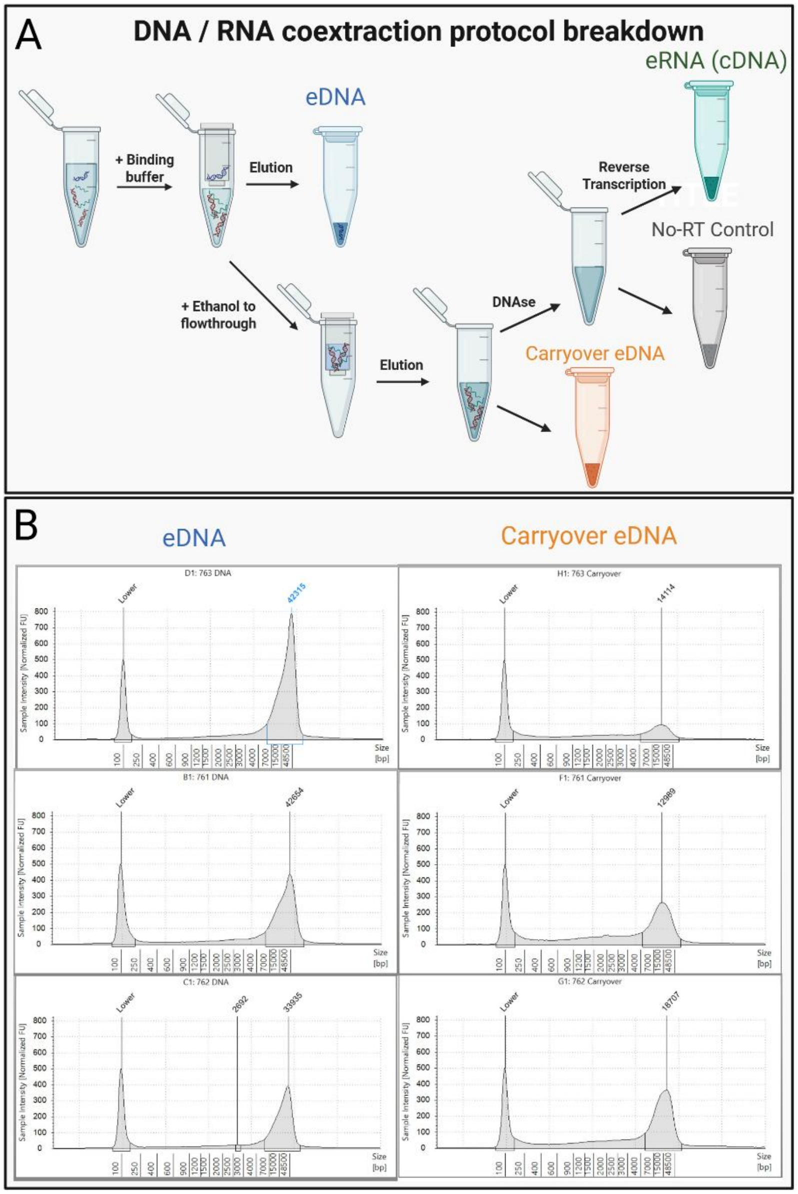

**Figure S2. Overview of DNA/RNA coextraction protocol and fragment size profiles of extracted DNA.** (A) Schematic representation of the coextraction workflow. After initial concentration using ultrafiltration and proteinase K digestion (not shown), binding buffer is added and the sample is passed through a first spin column. Environmental DNA (eDNA, blue) is eluted from this column following wash steps (not shown). The flowthrough is then mixed with ethanol to recover remaining nucleic acids and applied to a second spin column. The eluate from this second column—also obtained after washing—contains RNA along with residual DNA that did not bind to the first column, referred to as "carryover DNA" (orange). This second extract is split for downstream processing: one portion is treated with DNase to remove residual DNA prior to reverse transcription, while the other is retained untreated as the carryover DNA fraction. DNase-treated RNA is reverse transcribed into complementary DNA (eRNA/cDNA, green), and a parallel No-RT control (gray) is included to monitor for incomplete DNA removal. (B) Fragment analysis of eDNA and carryover DNA from three replicate samples. Left panels show electropherograms of eDNA, with dominant peaks at high fragment sizes. Right panels display the corresponding carryover DNA profiles prior to DNase treatment, showing lower-intensity, shorter DNA fragments, consistent with systematic fragmentation of DNA that escapes initial binding. Created in https://BioRender.com.

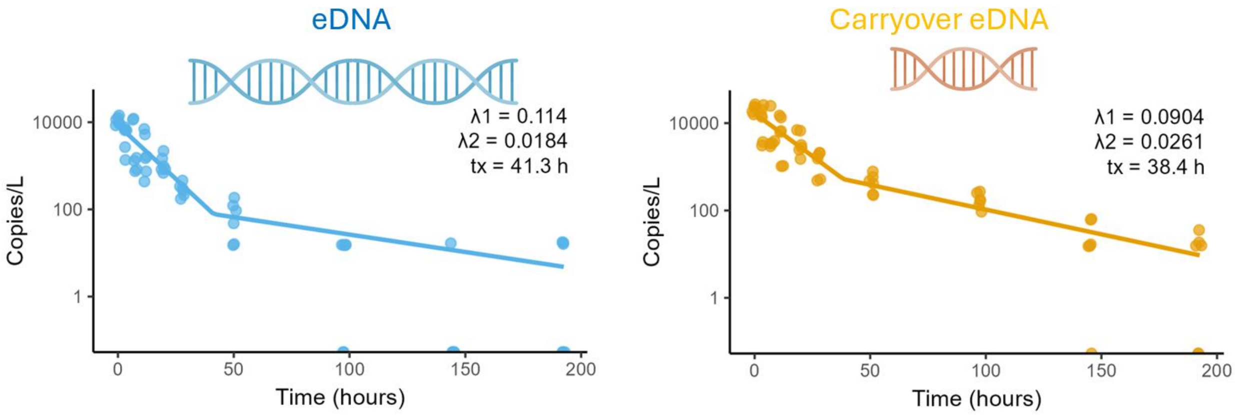

**Figure S3. Decay dynamics of eDNA and carryover eDNA over 188 hours.** Quantitative decay curves for mitochondrial *Cytb* marker in eDNA (left, blue) and carryover eDNA (right, orange). Both datasets were fitted with biphasic exponential decay models, capturing rapid initial loss ($\lambda_1$) followed by slower secondary degradation ($\lambda_2$), with the inflection point denoted as the transition time (tx). eDNA exhibits faster initial decay and slower long-term loss, while carryover DNA decays more gradually at first but stabilizes at a higher residual concentration, suggesting accumulation of short fragments or protection from degradation. Y-axis is on a log scale; points represent biological replicates.

### Metabarcoding of carboy water

The mitochondrial D-loop primers used for ddPCR amplify most odontocetes rather than being *Tursiops*-specific (Baker et al., 2018), and both killer whales (*Orcinus orca*) and harbor porpoise (*Phocoena phocoena*) inhabit the study area. Therefore, it was necessary to verify that our decay experiment was free of non-target cetacean DNA.

To do that, we opportunistically metabarcoded water collected at the initial (0 h) time-point from every carboy with two broad markers: (i) the same Dloop markers used in the main text (Table 1), and (ii) MiFish-U (Miya et al., 2015), a teleost marker that also amplifies marine mammals as by-catch. Dual-indexed Illumina-tailed primers were used in a two-step PCR workflow following Shaffer et al (in press). Libraries were pooled equimolarly and sequenced on an Illumina MiSeq (2 × 300 bp, v3 chemistry). Reads were denoised with DADA2 (Callahan et al., 2016), merged, chimera-filtered, and collapsed into amplicon-sequence variants (ASVs). Taxonomy was assigned with BLASTn against the NCBI nt database followed by least-common-ancestor filtering in TaxonKit (Shen & Ren, 2021).

Across both markers the only cetacean ASVs detected were assigned to the common bottlenose dolphin (*Tursiops truncatus*; Tables S2–S3), confirming that no DNA from *O. orca*, *P. phocoena* or other cetaceans was present, and that subsequent decay analyses exclusively track the target species.

**Table S2: Number of sequenced reads across species for the cetacean-specific Dloop marker.**

| Taxon | Class | Carboy 1 | Carboy 2 | Carboy 3 |
|---|---|---|---|---|
| ***Tursiops truncatus*** | **Mammalia** | **102087** | **150129** | **122654** |
| *Zalophus californianus* | Mammalia | 27 | 0 | 0 |
| NA | NA | 0 | 1277 | 66 |

**Table S3: Number of sequenced reads across species for the fish-specific MiFishU marker.**

| Taxon | Class | Carboy 1 | Carboy 2 | Carboy 3 |
|---|---|---|---|---|
| *Artedius fenestralis* | Actinopteri | 2181 | 1305 | 1913 |
| *Clupea pallasii* | Actinopteri | 35964 | 72788 | 45859 |
| *Cymatogaster aggregata* | Actinopteri | 0 | 4581 | 1361 |
| *Gadus sp.* | Actinopteri | 0 | 1203 | 0 |
| *Gasterosteus aculeatus* | Actinopteri | 787 | 821 | 2500 |
| *Leptocottus armatus* | Actinopteri | 0 | 0 | 19883 |

| | | | | |
|---|---|---|---|---|
| *Mallotus villosus* | Actinopteri | 5688 | 2828 | 2108 |
| *Merluccius productus* | Actinopteri | 0 | 493 | 0 |
| *Oligocottus sp.* | Actinopteri | 1556 | 0 | 0 |
| *Oncorhynchus keta* | Actinopteri | 712 | 0 | 3404 |
| *Pholis laeta* | Actinopteri | 0 | 8559 | 1484 |
| *Pholis ornata* | Actinopteri | 1543 | 0 | 3879 |
| *Homo sapiens* | Mammalia | 703 | 54 | 0 |
| *Phoca vitulina* | Mammalia | 0 | 501 | 1159 |
| ***Tursiops truncatus*** | **Mammalia** | **4232** | **1351** | **2317** |
| *Zalophus californianus* | Mammalia | 1095 | 2874 | 318 |
| **NA** | NA | 1026 | 4246 | 10108 |

### Alternative decay models

Before selecting the decay formula in our hierarchical model, we tested multiple alternative decay formulas commonly used in eDNA literature to determine the best-fitting approach for our data. The tested models included:

1. Exponential Decay, which is the standard single-phase exponential decay model:

$$\mathrm{C(t)} = \mathrm{C_0} e^{-\lambda \mathrm{t}}$$

   where $C(t)$ is the concentration of eNA at time t (copies/L), $\mathrm{C_0}$ is the initial concentration at t = 0 (copies/L), and λ is the decay rate constant (1/hour).

2. Biphasic Exponential Decay, which is a biphasic model that assumes two decay phases with a transition time:

$$\mathrm{C(t)} = \mathrm{C_0} e^{-\lambda_1 \mathrm{t}}\,, \;\; \mathrm{t} < \mathrm{t_x}$$

$$\mathrm{C(t)} = \mathrm{C_x} e^{-\lambda_2 (t - t_x)}\,, \mathrm{t} \geq \mathrm{t_x}$$

   Where $\lambda_1$ and $\lambda_2$ are the decay rate constants (per hour) for the second and first time phases, respectively, $\mathrm{t_x}$ is the time in hours where decay rate changes, and $\mathrm{C_x}$ is the concentration in copies/L at $\mathrm{t_x}$ determined as $\mathrm{C}_x = \mathrm{C_0} e^{-\lambda_1 \mathrm{t}_x}$

3. Power Law Decay, often used when decay slows at a constant rate over time:

$$\mathrm{C(t)} = \mathrm{C_0} \mathrm{t}^{-\lambda}$$

4. Time-Scaled Power Law Decay, which is a logarithmic power law model incorporating a time scale factor:

$$\mathrm{C(t)} = \mathrm{C_0} + \lambda \log(t\,.\,\alpha)$$

   Where α is a time scaling factor

5. Weibull Decay, which is a generalization of exponential decay with a variable rate across timepoints:

$$C(t) = C_0 e^{-\left(\frac{t}{\tau}\right)^{\alpha}}$$

Where $\tau = 1/\lambda$ if $\alpha = 0$

6. Logistic Decay, used in decay processes with saturation effects:

$$C(t) = \frac{C_0}{1 + e^{-\lambda(t - t_0)}}$$

Where $t_0$ is the inflection point with highest decay

We implemented six alternative versions of our joint ddPCR-decay model, each incorporating a different decay function. Model performance was evaluated using leave-one-out cross-validation (LOO-CV) by randomly leaving one sample out at a time via the LOO package in R (Vehtari et al., 2017).

Among the tested models, the biphasic exponential decay model provided the best fit to the data (Table S4). As a result, biphasic exponential decay was selected for all downstream analyses. We should also note that the Weibull model exhibited identifiability issues when estimating separate decay rates for each replicate, limiting its practical application.

**Table S4: decay model selection according to LOO-CV.**

| Decay Model | elpd_diff | se_diff |
|---|---|---|
| Biphasic Exponential | 0 | 0 |
| Weibull | -616.9 | 268.2 |
| Exponential | -1339.1 | 400.5 |
| Time Scaled Power | -2057.3 | 660.7 |
| Power Law | -3526.5 | 797.1 |
| Logistic | -5495.3 | 1071.0 |

### Raw data visualization

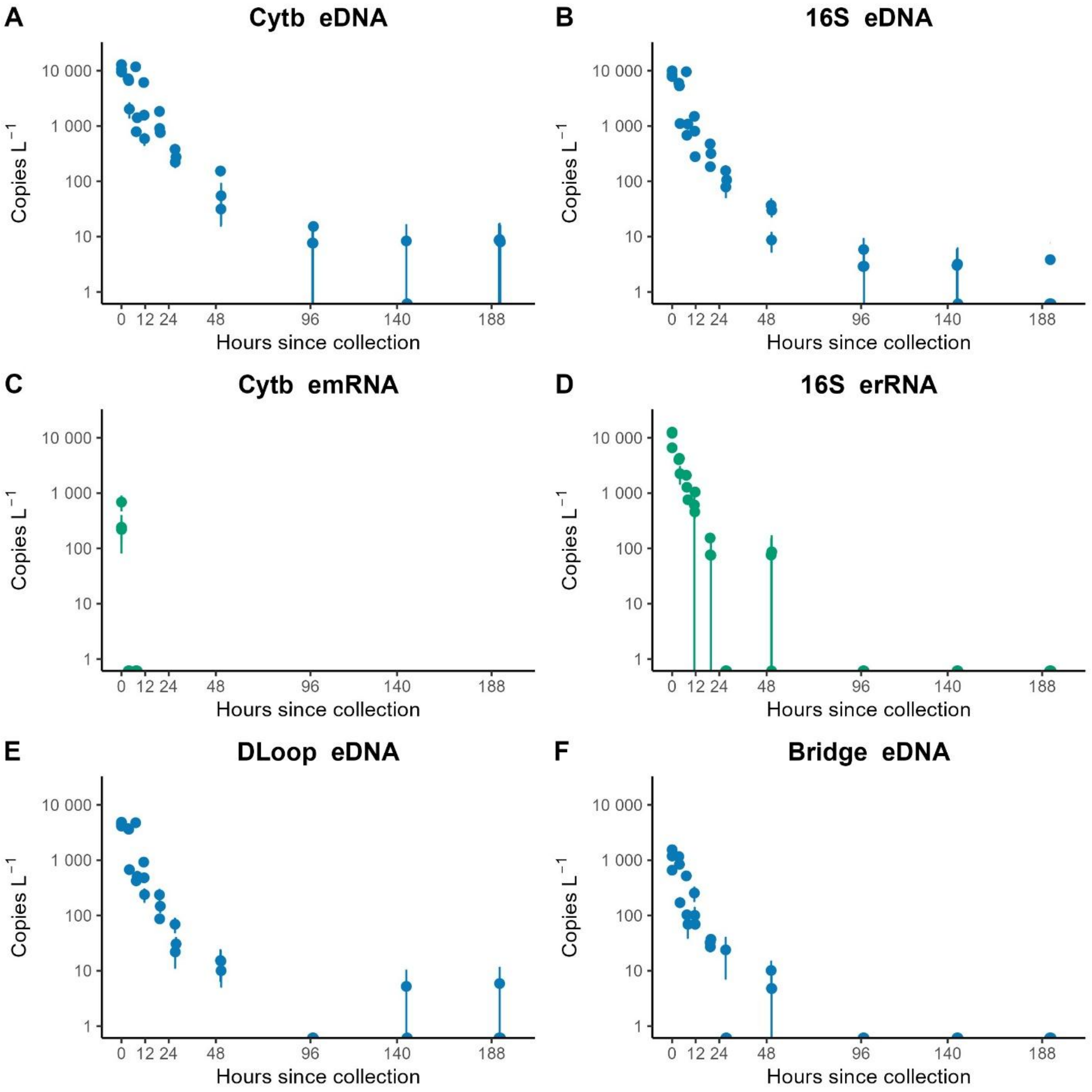

**Figure S4: Decay of environmental nucleic acids (eNAs) over time across multiple mitochondrial markers and components.** Each panel shows the concentration of dolphin-derived eNA in carboys (copies L$^{-1}$, log scale) over time since collection, based on ddPCR quantification of DNA (blue) and RNA (green) for six mitochondrial targets: (A) *Cytb* eDNA, (B) *16S* eDNA, (C) *Cytb* emRNA, (D) *16S* erRNA, (E) *DLoop* (DLL1) eDNA, and (F) a long-fragment "bridge" spanning 2,746 bp between *DLoop* and *16S*. Points represent means across technical replicates for each biological replicate (Carboy), with error bars showing ±1 SE. All measurements are from 5 μm filter fractions.
